# Passive transfer of fibromyalgia pain from patients to mice

**DOI:** 10.1101/713495

**Authors:** Andreas Goebel, Clive Gentry, Ulku Cuhadar, Emerson Krock, Nisha Vastani, Serena Sensi, Katalin Sandor, Alexandra Jurczak, Azar Baharpoor, Louisa Brieskorn, Carlos Morado Urbina, Angelika Sandstrom, Janette Tour, Diana Kadetoff, Eva Kosek, Stuart Bevan, Camilla I. Svensson, David A. Andersson

## Abstract

Fibromyalgia syndrome (FMS) is a chronic pain condition characterized by widespread pain and tenderness^1,2^. The etiology and pathophysiology of fibromyalgia are unknown and there are no effective treatments. Here we show that sensory hypersensitivity in FMS is caused by autoantibodies that act by sensitizing nociceptive sensory neurons. Administration of IgG from FMS patients increased mouse pain sensitivities to stimulation with mechanical pressure and cold. In contrast, transfer of IgG depleted samples from FMS patients or IgG from healthy control subjects had no effect on pain sensitivity. Sensory nerve fibres in *ex vivo* skin-nerve preparations from mice treated with FMS IgG were hypersensitive to mechanical stimulation. Immunohistochemical analysis revealed that IgG from FMS patients specifically labeled satellite glial cells and myelinated fibre tracts, as well as a small number of macrophages and endothelial cells in mouse dorsal root ganglia but not skin, muscle, spinal cord and brain. Our results demonstrate that fibromyalgia pain is caused by IgG autoantibodies that sensitize peripheral nociceptive afferents neurons and suggest that therapies that reduce patient IgG titres may be effective treatments of fibromyalgia pain.

## MAIN

Fibromyalgia syndrome (FMS) is a chronic pain condition characterized by widespread pain, augmented pain sensitivity to mechanical pressure and cold temperatures^3–5^, as well as fatigue and emotional distress^6–8^. The prevalence of FMS is approximately 2%^9^, and at least 80% of FMS patients are women. The prevalence rises to 10-30% among patients diagnosed with autoimmune rheumatological conditions^10,11^, and FMS is thus one of the most common chronic pain conditions. The etiology and pathophysiology of FMS are unknown and the current treatment strategies for FMS rely mainly on life style changes, physical exercise and drug therapy with antidepressants and anticonvulsants^1^. Unfortunately, the modest efficacy of the available therapies in most patients leaves an enormous unmet clinical need. The animal models that have been used for experimental studies of FMS have uncertain translational relevance, and rely on local repeated intramuscular injections of acid^12^ or systemic depletion of monoamines by reserpine treatment^13^. Consequently, the development of novel, mechanism-based therapies has been hampered by the limited understanding of the basis of FMS.

The increased polymodal pain sensitivity experienced by FMS patients^14,15^ is associated with altered pain processing in the central nervous system^2^, dysfunctional descending pain modulation^16,17^, and structural and functional changes in the brain^18–20^. FMS is also associated with abnormalities in peripheral sensory afferents, such as spontaneous ectopic activity and sensitization of C-fibers, and loss of epidermal innervation^21,22^. Furthermore, elevated levels of proinflammatory cytokines^23,24^, and reduced levels of anti-inflammatory cytokines^25,26^ have been detected in FMS patients, suggesting that inflammatory processes may be engaged. These observations, together with the markedly increased prevalence of FMS among patients with autoimmune rheumatological conditions^10,11^, led us to hypothesize that FMS may have an autoimmune basis. Here we have investigated the possibility that IgG autoantibodies cause FMS pain by examining whether sensory abnormalities can be transferred to mice by administration of IgG purified from FMS patients.

We administered IgG purified from the serum of individual FMS patients and healthy control (HC) subjects to female mice by intraperitoneal injection for 4 consecutive days (8mg per day), a dose regimen based on the original studies identifying myasthenia gravis as an autoantibody mediated disorder^27,28^. Since FMS patients regularly report pressure hypersensitivity, we examined paw withdrawal thresholds in mice using the Randall-Selitto paw-pressure test following IgG transfer. IgG from each of the 8 patients (supplementary table 1), but not from any of the 6 healthy control subjects, rapidly produced mechanical hypersensitivity (Fig. 1a, e, g). In addition to pressure sensitivity, patients frequently report that pain is exacerbated by cold temperatures, and quantitative sensory testing has demonstrated an increased cold pain sensitivity in FMS^4,15^. In good agreement with patient observations, administration of IgG from FMS patients gave rise to a significantly increased sensitivity to noxious cold in mice (Fig.1b, f, h). Both mechanical and cold hypersensitivities were established within 24-48 hours after the first injection and maintained for at least about a week after the final IgG injection. FMS IgG also generated hypersensitivity to stimulation with calibrated von Frey filaments, a widely used test of mechanical nociception in mice (Fig. 1c). Fibromyalgia pain is widespread, and we therefore examined the pressure sensitivity of the thigh (Fig. 1d) using the Randall-Selitto device, to determine whether FMS IgG affects the pressure sensitivity of sites other than the hind paw in mice. In this test, FMS IgG produced significant mechanical hypersensitivity in the thigh compared to treatment with IgG from HC. The observed hypersensitivities produced by IgG preparations from different FMS patients showed similar amplitudes (Fig. 1e, f) and time courses, which is reflected in the plot showing the averaged mechanical and cold sensitivities produced by IgG from 8 individual FMS patients and 6 HC subjects over 10 days (Fig. 1g, h). These results demonstrate that the transfer of sensitivities from FMS subjects to mice is reproducible across multiple FMS subject, strongly suggesting that an antibody-dependent mechanism may underlie FMS pain.

**Figure 1.**
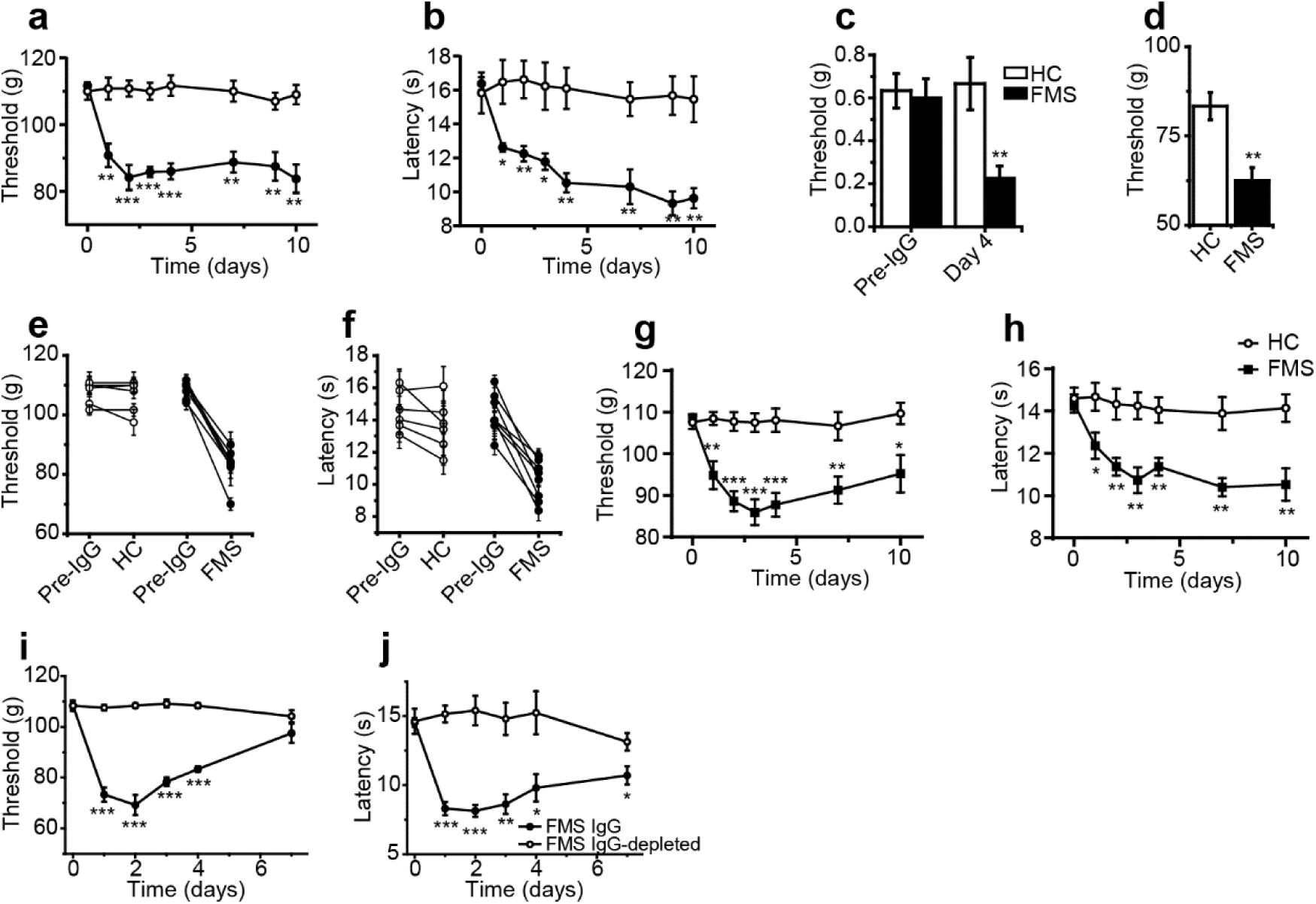
Passive transfer of hypersensitivities from fibromyalgia patients to mice. Administration of IgG from an FMS patient reduced the paw withdrawal threshold in the paw-pressure test (**a**), the withdrawal latency in the cold-plate test (**b**), the response threshold to stimulation with von Frey filaments (**c**), and the withdrawal threshold to mechanical pressure of the thigh (**d**), compared to administration of IgG from healthy control subjects. Administration of purified IgG from each of 8 FMS patients evoked mechanical (**e;** p between 0.02 and 2×10^-5^) and cold hypersensitivity (**f**, p between 0.05 and 2×10^-4^), whereas 6 preparations from HC did not (all p>0.25 in the paw pressure test and p>0.15 in the cold plate test). Paw withdrawal threshold or latency before injection of IgG and at the time of maximum sensitivity during the 7 days following IgG injections are shown in **e, f**. Mean sensitivity in the paw-pressure test (**g**) and cold plate test (**h**) following administration of IgG from 8 patients and 6 control subjects (each experiment performed as in **a, b)**. IgG from a single patient, but not the IgG-depleted serum from the same patient, produced mechanical (**i**) and cold (**j**) hypersensitivity. Data points are mean ± SEM of n=6 mice in **a-d**. Data points in **e**-**f** are mean ± SEM of 8 (FMS) or 6 (HC) experiments, each performed with n=4-6 mice per treatment. *P<0.05, **P<0.01, ***P<0.001, FMS IgG compared to HC IgG, unpaired two-tailed t-test.

We next compared the effects of FM patient IgG with IgG-depleted patient serum to assess whether immunoglobulins and serum components other than IgG may contribute to pain and hypersensitivity in patients (Fig.1i, l). As expected, IgG from a single patient produced marked cold and mechanical hypersensitivities, whereas the IgG-depleted serum from the same patient was without any effect (Fig.1i, j).

To determine whether autoantibodies are responsible for pain and painful hypersensitivities in a wider and regionally distinct cohort of FMS patients, we examined the effects of transferring IgG preparations pooled from 20 FMS patients and 20 healthy control subjects (Supplementary table 1). As expected, FMS patients displayed markedly higher pain ratings on a visual-analogue scale (VAS; Fig. 2a) and a significantly increased pressure pain sensitivity compared to healthy control subjects (Fig. 2b). Administration of pooled IgG to mice (two separate experiments, each using IgG pooled from separate groups of 10 patients and 10 control subjects) significantly increased the sensitivity in the paw-pressure test compared to administration of pooled IgG from control subjects (Fig. 2c). In these experiments we used the same dose regimen as above (8mg of IgG on four consecutive days), but since the IgG from each individual donor was diluted 10-fold, the observed pro-nociceptive effect demonstrates that IgG from different patients have additive activity. Similar to our observations with IgG from individual patients, the pooled FMS IgG also elicited hypersensitivity to cold (Fig. 2d) and to stimulation with von Frey filaments (Fig. 2e). To determine the duration of the IgG-mediated hypersensitivity we studied mechanical thresholds in mice treated with IgG pooled from patients or control subjects for one month. The onset of mechanical hypersensitivity produced by pooled FMS IgG followed the same time course as seen with IgG from individual donors and resolved fully about 2.5 weeks after cessation of IgG administration (Fig. 2f). This observation indicates that the painful hypersensitivities produced by FMS IgG are reversible and that sensory abnormalities resolve when the levels of pathogenic IgG are reduced sufficiently. Our results thus strongly suggest that therapies which reduce patient IgG levels may be effective treatments for FMS pain.

**Figure 2.**
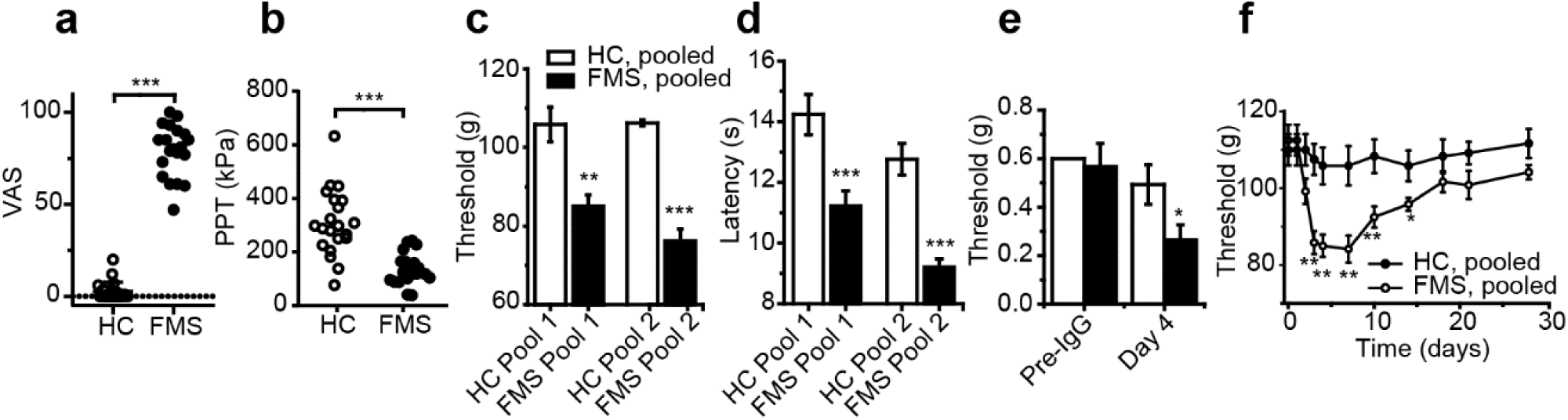
Passive transfer of hypersensitivity by pooled IgG. Visual analogue pain scores (**a**) and pressure pain thresholds (**b**) reported by 20 FMS patients and 20 healthy control subjects. Administration of IgG from 2 pools of 10 FMS patients produced mechanical (**c**), cold (**d**) and tactile (von Frey, pool 1, **e)** hypersensitivity in mice compared to 2 pools of HC IgG (each pool from 10 subjects). Time course of mechanical hypersensitivity following administration of IgG pooled from 10 FMS patients and HC (**f**). *P<0.05, **P<0.01, ***P<0.001, FMS IgG compared to HC IgG, unpaired two-tailed t-test or Mann-Whitney U-test (**a, b**) or unpaired two-tailed t-test (**c**-**f**).

### Sensitization of nociceptors

Microneurography studies of FMS patients have demonstrated sensitization and ectopic impulse discharge in nociceptive C-fibres^21^. To determine whether the sensory abnormalities demonstrated in mice *in vivo* following administration of FMS patient IgG could be explained by functional modification of peripheral nociceptors, we next examined the electrophysiological properties of single afferent nerve fibre units in skin-saphenous nerve preparations from IgG treated mice. Single units were classified according to their conduction velocity and mechanical response threshold, as described previously^29,30^.

We stimulated the receptive fields of Aδ- and C-mechanosensitive nociceptors (AM and CM fibers) mechanically and compared the responses in preparations from mice after 4 days of treatment with IgG from either HC or FMS patients (Fig. 3). Both AM and CM fibers in preparations from mice treated with FMS IgG responded to mechanical stimulation with a significantly increased number and frequency of action potentials compared to preparations from HC IgG treated mice (Fig. 3a-d). The increased impulse activity evoked by mechanical stimulation is consistent with the C-fibre sensitization observed in FMS patients using microneurography^21^ and demonstrates that patient autoantibodies produce a heightened nociceptor activity that is maintained in the absence of the central nervous system. Since we had observed a robust behavioral cold hypersensitivity in mice treated with FMS IgG, we next examined whether this phenotype was accompanied by functional abnormalities in cold and mechanosensitive Aδ-fibres (AMCs). We used cold superfusion to identify the receptive fields of AMCs and thereafter challenged these with a 60s cooling ramp (from 32°C to 5-9°C). Cooling evoked a significantly increased number of action potentials in AMCs in preparations from mice treated with FMS IgG compared to HC IgG (Fig.3 e, f; FMS median 82, interquartile range 41-353; HC 22, 3-52), whereas as the cold activation thresholds did not differ significantly (Fig.3 g). Analysis of the temporal impulse pattern demonstrated that AMCs from FMS IgG treated mice discharged action potentials at a higher rate throughout the cooling ramp, consistent with a heightened excitability (Fig.3 h).

**Figure 3.**
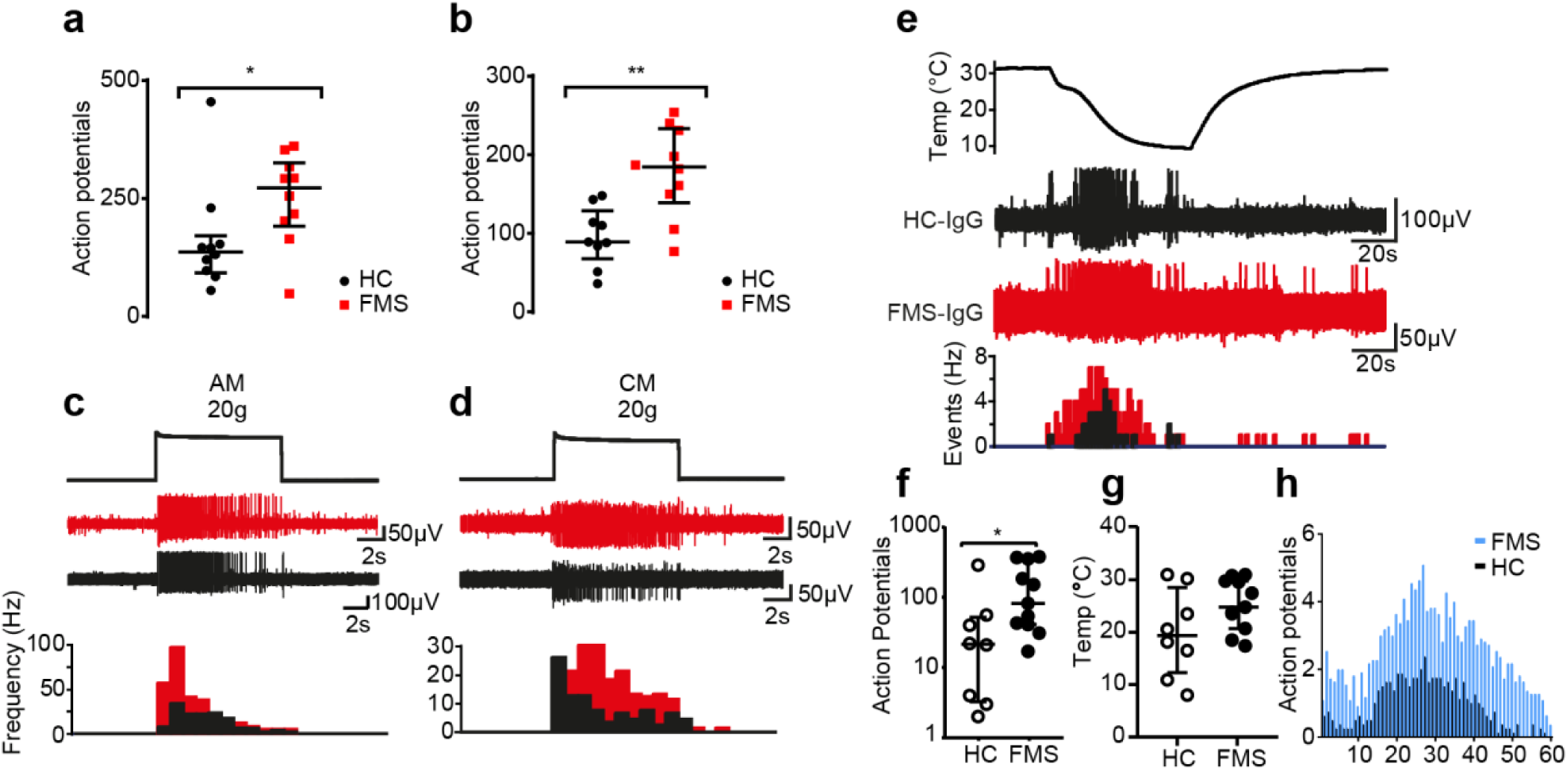
FMS IgG sensitizes nociceptors. The number of action potentials elicited by a force step (20g, 10s) was significantly increased in Aδ- (**a**) and C-mechanonociceptors (**b**) in preparations from mice treated with FMS IgG compared to HC IgG. Typical recordings of Aδ- (“AM”) and C-mechanonociceptors (“CM”) in FMS and HC preparations (**c-d)**. The lower panels show the increased action potential frequency in FMS IgG treated preparations compared to HC IgG. Typical recordings of cold-sensitive AM fibres (AMCs) in preparations from mice treated with FMS or HC IgG (**e**). The number of cooling-evoked action potentials (**f**) the temperature activation thresholds (**g**) in AMCs from mice treated with FMS IgG compared to HC IgG. The average impulse pattern during 60s cooling ramps demonstrate an increased discharge rate in AMCs from FMS IgG treated mice. Line and whiskers indicate median and interquartile range. *P<0.05, **P<0.01, FMS IgG compared to HC IgG, Mann-Whitney U-test.

**Figure 4.**
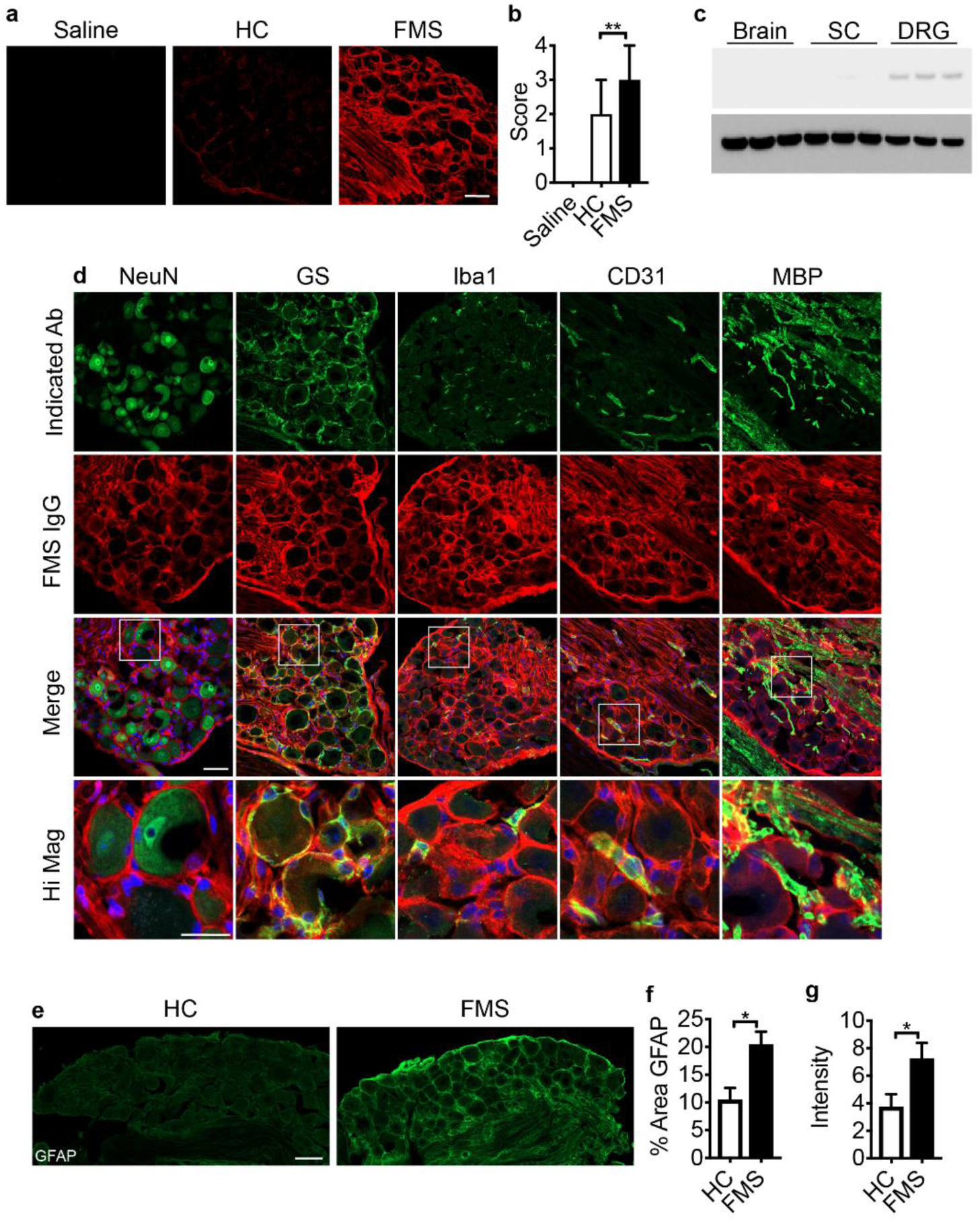
FMS IgG accumulates in the DRG and binds satellite glial cells. FMS IgG, but not HC IgG accumulates in the dorsal root ganglia (DRG) 14 days after the first IgG injection (**a, b**). Western blot analysis detected FMS IgG in the DRG but little to no IgG in spinal cords (SC) or brains (**c**) at the last day of IgG injection (day 4). FMS IgG immunoreactivity does not colocalize with neurons (NeuN positive cells) but does colocalize with satellite cells (glutamate synthase (GS) expressing cells), some macrophages (Iba1 expressing cells) and blood vessels (CD31 expressing cells) and myelinated fibre tracts (myelin basic protein (MBP) staining) within but not outside the DRG compartment (**d**). DRGs from FMS IgG injected mice have increased GFAP immunoreactivity, which is indicative of increased satellite glial cell activity, compared to HC injected mice (**e**) when the % area of GFAP immunoreactivity (**f**) and GFAP mean pixel intensity (g) are quantified. Human IgG immunoreactivity was quantified using a custom 0 (no IgG) to 4 (very strong IgG signal), n=9-10, data points are presented as median ± 95% C.I. GFAP data points are mean ± SEM, n=5-6. * indicates p<0.05, ** indicates p<0.01 compared to HC IgG, Kruskal-Wallis test followed by a Mann-Whitney test (b) or t-test (**f, g**). Scale bars indicate 50 μm, except the high magnification image scale bar which indicates 25 μm.

### Immunohistochemical localization of human IgG

Immunohistochemical analysis of tissues from mice that had been injected with FMS IgG using anti-human IgG antibodies revealed robust staining in dorsal root ganglia (DRG; Fig. 3a), while no specific immunoreactivity was observed in the spinal cord (Fig.3 c, Supp. Fig. 1a). In contrast, IgG from control subjects generated only low levels of immunoreactivity in DRGs (Fig. 3a, b). FMS IgG was also reliably detected in DRG, but neither in brain nor spinal cord tissue by western blot analysis (Fig. 3c). The FMS patient IgG was primarily located to satellite glial cells (visualized by glutamine synthase) and to fibre tracts entering the DRG, as well as a number of Iba1 positive macrophages and CD31 positive blood vessels (Fig 3d). FMS patient IgG did not colocalize with DRG neurons (visualized by NeuN). In addition to the specific localization of FMS IgG to satellite glial cells, we noted signs of enhanced satellite cell activity, based on changes in GFAP immunoreactivity, in mice injected with FMS IgG compared to HC IgG injected controls (Fig 3e-g). In contrast, no changes in astrocyte (GFAP; Supp. Fig. 1b-d) or microglia (Iba1; Supp. Fig. 1e-g) reactivity were observed in the spinal cord. In combination with the lack of FMS IgG in the brain of spinal cord, these observations strongly suggest that the pronociceptive actions of FMS IgG were peripheral. Future investigations will determine whether an action of IgG on satellite cells is responsible for FMS pain and elucidate the molecular mechanisms underlying the neuronal hypersensitivities. It is clear, however, that satellite glial cells and sensory neurons can be electrically coupled and that satellite cells can influence neuronal excitability^31–33^.

Here we identify FMS as an autoimmune pathology in which IgG purified from FMS patients evokes painful hypersensitivity by sensitizing peripheral nociceptors when transferred to mice. A peripheral origin of pain and tenderness agrees well with earlier reports of peripheral sensory abnormalities in patients^5,15,21,22^. A chronically heightened noxious peripheral input may be expected to generate altered patterns of activity in the CNS^20^. Identification of FMS as an autoimmune disease, rather than an unexplained functional somatic syndrome, will transform future fibromyalgia research and facilitate development of mechanism-based therapeutic interventions. Our results suggest that therapies that reduce the IgG titre in patients, or which prevent the functional consequences of the pathological IgG, will be effective therapies for fibromyalgia.

## Acknowledgements

This work was supported by the Medical Research Council MR/S003428/1 (DAA, SB and AG), Versus Arthritis 21544 (DAA and AG) and 21543 (SB) the Pain Relief Foundation (DAA and AG), the Swedish Research Council (CIS), Knut and Alice Wallenberg Foundation (CIS), the Karolinska Institute Foundations (CMU), the EU Projects FP7-Health-2013-Innovation-1602919-2 (CIS), European Union Seventh Framework Programme (FP7/2007–2013) under grant agreement no 602919, the IASP John J. Bonica fellowship (EK) and by a generous donation from Leif Lundblad and family (CIS, EK).

---

**Supplemental Figure 1.**
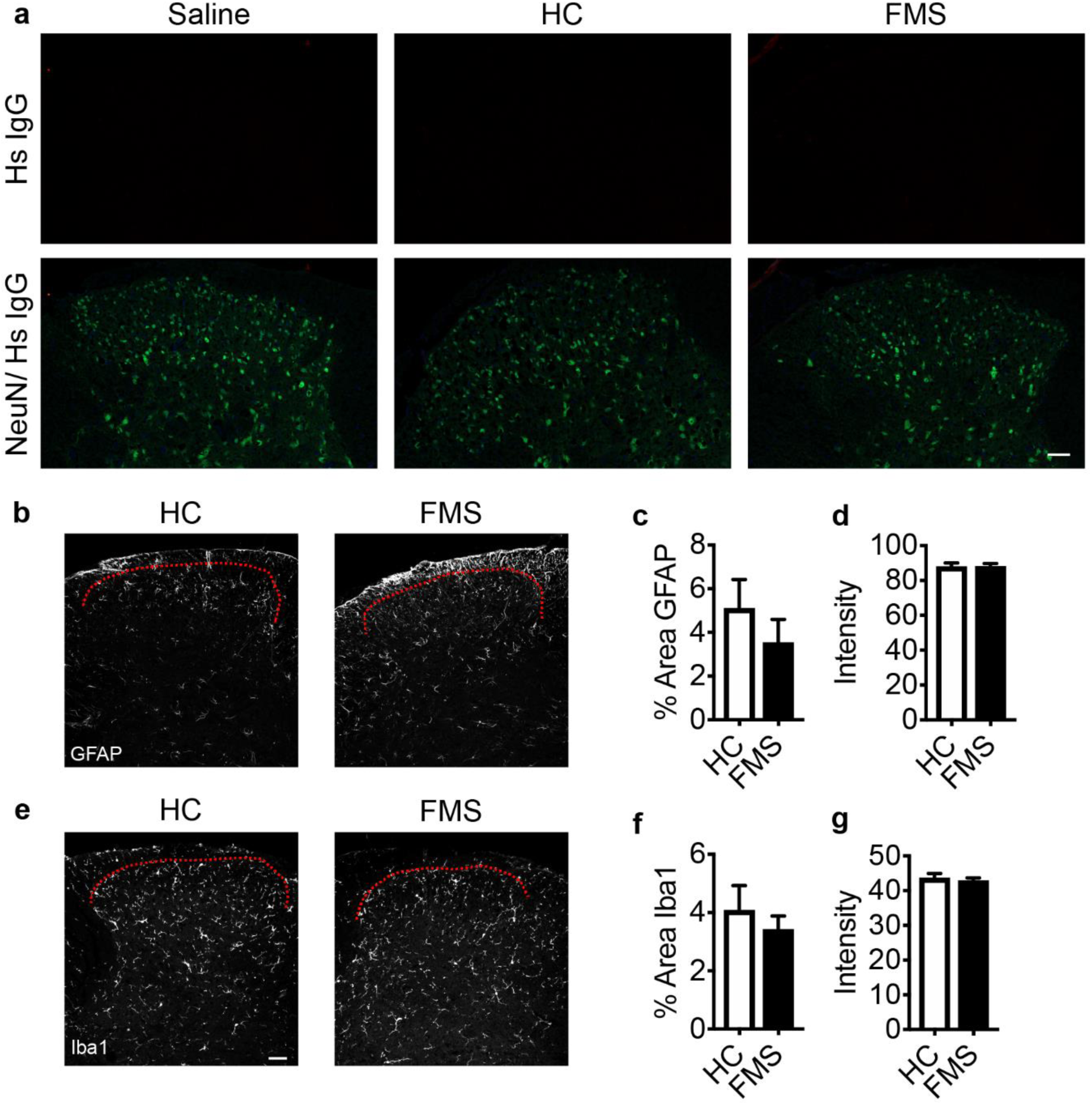
FMS IgG does not accumulate in the spinal cord or activate dorsal horn glia. Neither HC IgG nor FMS IgG accumulates in the dorsal horn of the spinal cord (**a**). Sections were probed with an anti-Hs (human) IgG antibody (top row) and anti-NeuN antibody (bottom row, merged with anti-Hs IgG, **a**). FMS IgG did not change the percent area of GFAP immunoreactivity of GFAP intensity in the dorsal horn compared to HC IgG (**b-d**), suggesting that FMS IgG does not alter astrocyte activity. Iba1 percent area of immunoreactivity and intensity were also unchanged by FMS IgG compared to HC IgG (**f-g**), suggesting that FMS IgG does not alter microglia activity. Red dotted lines indicate dorsal edge of lamina I of the dorsal horn. Data points are mean ± SEM, n=5-6. Scale bars are 50 μm.

## METHODS

### Patient samples

Serum samples for individual testing were derived from UK patients managed for their Fibromyalgia at a department of pain medicine, or from healthy controls (Ethics North-West Haydock, Ref: 15/NW/0467). Pooled samples were obtained from Swedish patients or healthy controls responding to a study advert (approved by the local ethical committee 2011/2036–31/1). Individual informed consent was obtained from all patients. All patients had been examined by a Consultant rheumatologist, and UK patients had additionally been examined by a Consultant in pain medicine. We purified serum-IgG from 28 FMS patients and 26 healthy control subjects (HC). All patients fulfilled both 1990 and 2011 ACR diagnostic criteria for fibromyalgia^1,2^. Most patients (26/28) and all of the pooled sample donors were unaffected by other sensory or rheumatological conditions. Most patient donors (27/28) were women. The donors’ demographics and disease characteristics are provided in Supplemental tables 1-2.

**Supplemental table 1.** Patient characteristics. Age - mean (range), duration of FMS pain – median (interquartile range); pain intensity - average pain intensity over the past week ± SD (range); UK sample – numeric rating scale pain score on an 11-point scale (0-10), Swedish sample – visual analogue scale pain score (0-100), with 10, 100 = ‘worst pain imaginable’; tender point count – average (range) number of defined points which are tender to pressure out of a maximal 18.^1^

|  | Age; %female | Duration (years) | Pain intensity | Tender point count |
| --- | --- | --- | --- | --- |
| UK sample, n=8 | 46 (19-58); 88 | 2.5 (1.3-18) | 8.1±0.8 (6.5-9) | 15 (12-18) |
| Swedish sample, n=20 | 49 (35-58); 100 | 11 (3-15) | 79±14 (47-100) | 17 (11-18) |

### Purification of immunoglobulins

IgG for individual testing was purified as described previously^3^, using protein G beads (Sigma-Aldrich, Gillingham, UK). Serum was diluted 1:3 with Hartmann’s solution, passed through a protein G column, and the bound IgG was eluted using 100 mM glycine pH 2.3; the pH was adjusted to 7.4 using 1 M Tris pH 8 and then dialyzed overnight at 4°C in Hartmann’s using a 10 kDa dialysis membrane (Fisher Scientific, Loughborough, UK). The concentration of IgG present after dialysis was determined using a modified Lowry assay (DC protein assay, BioRad, Hemel Hempstead, UK) and adjusted by dilution with Hartmann’s or by dialysis against a sucrose solution (Sigma-Aldrich). Finally, the IgG solution was sterile filtered using syringe-driven 0.2 µm filter units (Millipore, Watford, UK), stored at 4°C and used within 3 months. Sera for pooled sample testing was purified using HiTrap Protein G HP columns (GE Healthcare), eluted with 0.1M glycine:HCL pH2.7, and the pH adjusted to 7.4 with 1M Tris pH 9; consequently samples were dialysed against PBS, concentration adjusted, stored at −20C and later thawed, pooled, and concentration-adapted using PBS pre-wet concentration columns (Pall Corporation Macrosep 10K); concentration measurements were always with Nanodrop 2000 (Fisher Scientific).

### Animals

Behavioral experiments were carried out according to the U.K. Home Office Animal Procedures (1986) Act. All procedures were approved by the King’s College London Animal Welfare and Ethical Review Body and conducted under a UK Home Office Project License. Behavioural experiments were performed on female C57Bl/6J mice (8–10 weeks old) obtained from Envigo UK Ltd., Bicester, UK, housed in a temperature-controlled environment with a 12h light/dark cycle with access to food and water *ad libitum*. Immunofluorescence experiments were performed on 4-month old female BALB/c mice (Janvier, France) and were approved by the local ethical committee (4945-2018). Mice were injected intraperitoneally with 8 mg of IgG from healthy control subjects or FMS patients on 4 consecutive days.

### Behavioral studies

Before any nociceptive testing, mice were kept in their holding cages to acclimatize (10-15 min) to the experimental room. Mice were randomized between cages and the experimenter blinded to their treatment.

The Randall-Selitto paw-pressure test was performed using an Analgesymeter (Ugo-Basile, Italy). The experimenter lightly restrained the mouse and applied a constantly increasing pressure stimulus to the dorsal surface of the hind paw using a blunt conical probe. The nociceptive threshold was defined as the force in grams at which the mouse withdrew its paw^4^. A force cut-off value of 150 g was used to avoid tissue injury.

Tactile sensitivity was assessed using von Frey filaments (0.008-2 g) according to Chaplan’s up-down method^5^. Animals were placed in a Perspex chamber with a metal grid floor allowing access to their plantar surface and allowed to acclimatize prior to the start of the experiment. The von Frey filaments were applied to the plantar surface of the hind paw with enough force to allow the filament to bend, and held static for approximately 2-3 s. The stimulus was repeated up to 5 times at intervals of several seconds, allowing for resolution of any behavioral responses to previous stimuli. A positive response was noted if the paw was sharply withdrawn in response to filament application or if the mouse flinched upon removal of the filament. Any movement of the mouse, such as walking or grooming, was deemed an unclear response, and in such cases the stimulus was repeated. If no response was noted a higher force hair was tested and the filament producing a positive response recorded as the threshold.

Thermal sensitivity was assessed using a cold-plate (Ugo Basile, Milan). Paw withdrawal latencies were determined with the plate set at 10 °C. The animals were lightly restrained (scruffed) and the left hind paw was placed onto the surface of the plate^6,7^. The latency to withdrawal of the paw was recorded as the endpoint. A maximum cut-off of 30 seconds was used for each paw.

### Skin-nerve recording

Mice were killed by cervical dislocation and the hind paw was shaved prior to dissection of the isolated skin-nerve preparation. The saphenous nerve and the shaved skin of the hind limb were placed in a recording chamber at 32 °C. The chamber was perfused with a gassed (95% O_2_ and 5% CO_2_) prewarmed synthetic interstitial fluid (SIF): 108mM NaCl, 3.5mM KCl, 0.7mM MgSO_4_, 26.2mM NaCO_3_, 1.65mM NaH_2_PO_4_, 1.53mM CaCl_2_, 9.6mM sodium gluconate, 5.55mM glucose and 7.6mM sucrose. The skin was placed inside up (corium side up) and pinned down using insect pins (0.2 mm diameter) in the organ bath to allow access to the receptive fields. The saphenous nerve was placed through a small gap from the organ bath to an adjacent recording chamber on a mirror platform. The desheathed saphenous nerve was covered with paraffin oil for electrical isolation and dissected fine nerve filaments were placed on a fine gold wire-recording electrode using a microscope^8,9^.

### Conduction velocity

The saphenous nerve was divided into progressively thinner filaments until a single unit could be isolated in response to stimulation of the receptive field with a glass rod. The electrical latency of identified units (Digitimer DS2, Digitimer Ltd, Welwyn Garden City, UK) was used to determine the conduction velocity and to categorize units as Aβ (velocity>10m/s), Aδ (1.2<velocity<10m/s) or C-fibers (0<velocity<1.2m/s).

### Mechanical stimulation

A computer-controlled stimulating probe, equipped with a force transducer, was used to deliver mechanical stimuli to the most sensitive point of a receptive field (Avere Solutions UG, Erlangen, Germany). The number of action potentials elicited by stimulation with a 10s force step of 20g was determined in A- and C-mechanonociceptors (AM and CM fibers) in preparations from mice treated with FMS or HC IgG. Recording and analysis were done using Spike 2 (Cambridge Electronic Design, Cambridge, UK).

### Immunofluorescence

Mice were deeply anesthetized with isoflurane and perfused with PBS, followed by 4% PFA. The lumbar spinal cord and lumbar dorsal root ganglia were collected, post fixed for 24 hours in 4% PFA, cryoprotected in 20% sucrose and then embedded in optimal cutting temperature (OCT) compound (Sakura Finetek). Tissue was sectioned with a CryoStar NX70 cryostat (Thermo Scientific) at a thickness of 10 μm and thaw mounted onto SuperFrost Plus slides (Thermo Scientific) and slides were stored at −20°C until use. Prior to use slides were thawed for at least 30 minutes at room temperature and washed in PBS to remove excess OCT. Slides were blocked with PBS supplemented with 0.3% TritonX-100 and 3% normal goat or donkey serum, depending on the secondary antibody and then incubated with primary antibodies diluted in PBS supplemented with 0.1% Triton X-100 and 1% normal goat or donkey serum. Slides were washed and incubated with appropriate secondary antibodies (indicated in supplementary table 2). For colocalization studies of human IgG with various cell-type markers the primary antibodies against the cell type markers were first added and then following washing the anti-human IgG antibody was co-incubated with the secondary antibodies against cell-type markers. Following washing, the slides were incubated with DAPI (1:20000), washed again and then cover slipped with Prolong Gold mounting media (Thermo Scientific).

**Supplemental table 2.** Antibodies used.

| Primary Antibodies |  |  |  |  |  |
| --- | --- | --- | --- | --- | --- |
| Target | Host | Manufacturer | Catalogue # | Dilution |  |
| GS | Rabbit | Abcam | ab73593 | 1:500 |  |
| GFAP | Mouse | Millipore | Mab360 | 1:500 (DRG)<br>1:1000 (SC) |  |
| Iba1 | Rabbit | Wako | 019-19741 | 1:500 (DRG)<br>1:1000 (SC) |  |
| NeuN conj.<br>AF488 | Mouse | Millipore | MAB377X | 1:100 |  |
| MBP | Rat | Abcam | 550274 | 1:500 |  |
| CD31 | Rat | BD Pharmigen | Ab739 | 1:500 |  |
| Secondary Antibodies |  |  |  |  |  |
| Target | Fluorophore | Host | Manufacturer | Catalogue # | Dilution |
| Rabbit IgG | AF488 | Goat | ThermoFisher | A11008 | 1:250 |
| Human IgG | AF594 | Goat | ThermoFisher | A11014 | 1:200 |
| Mouse IgG | AF488 | Goat | ThermoFisher | A11029 | 1:250 |
| Rat IgG | AF488 | Goat | ThermoFisher | A11006 | 1:250 |

GS (glutamate synthase), GFAP (Glial fibrillary acidic protein), NeuN (Neuronal N), MBP (Myelin Basic Protein), AF (Alexa Fluor)

### Fluorescent microscopy and analysis

DRGs were imaged using a confocal microscope (Zeiss LSM800) operated by LSM ZEN2012 (Zeiss) software. For each animal 5 DRG sections separated at least 50 μm were analyzed and averaged. Human IgG accumulation was assessed by two blinded scorers using a 0-4 visual scoring system and the GFAP signal was analyzed in ImageJ as % GFAP positive area and average GFAP pixel intensity. Spinal cords were imaged using a Nikon Eclipse TE300 fluorescent microscope with a Nikon Digital Sight DS-Fi1 camera. Iba1 and GFAP signal intensity were analysed by two blinded reviewers using a custom developed Python script. For each animal 5 spinal cord sections separated at least 50 μm were analyzed and averaged and the data presented as mean ± SEM. Representative spinal cord images were taken using a Zeiss LSM 800 confocal microscope.

### Western blot analysis

Mice were deeply anesthetized with isoflurane and euthanized by decapitation. Brain, spinal cord and lumbar DRG were collected and flash frozen for later use. Protein was extracted from the tissue by sonication in an extraction buffer (0.5% Triton X-100, 50mM Tris, 150 mM NaCl, 1mM EDTA and 1% SDS in water, pH 7.4). Protein concentration was analyzed using with the BCA Assay (Pierce). Samples were diluted in an LDS loading buffer (NuPage) and denatured with DTT at 95°C. An equal amount of protein per sample was loaded into 2-12% gradient NuPage Bis-Tris gels (Thermo Scientific) and protein was separated by electrophoresis. Protein was transferred to a nitrocellulose membrane using a dry transfer system (Thermo Scientific). The membrane was blocked with 5% non-fat milk powder (Biorad) in TBS.T for 1 hour at room temperature. The membrane was then probed with an anti-human HRP conjugated antibody (Santa Cruz) diluted in TBS.T with 5% non-fat milk powder. SuperSignal West Pico Chemiluminescent Substrate (Thermo Scientific) was incubated with the membrane and protein was visualized using a Biorad ChemiDoc™ Image System.

### Reagents

All buffers and salts were purchased from Sigma (Poole, UK), VWR (Lutterworth, UK) or Thermofisher (Waltham, MA, USA).

### Statistics

Data are presented as individual data points or as mean±SEM and the number of animals or single units studied are indicated by n. No data points have been removed from any of the datasets presented. Data from behavioral *in vivo* experiments were tested statistically by unpaired, two-tailed t-test, whereas patient data and electrophysiological data were tested by Mann-Whitney U-test.

